# Trait-space patterning is dictated by the tempo and mode of mutation

**DOI:** 10.1101/2024.11.21.624779

**Authors:** Stephen Martis, David J. Schwab, Trevor GrandPre

## Abstract

In large, natural ecosystems, many (≳ 1) phenotypically relevant mutants can emerge over the characteristic turnover time of the population. When this is the case, there can be ‘eco-evolutionary feedback’ between the dynamical processes that underlie mutation, selection and ecology. In this paper, we show that, owing to such feedback, the precise details of the mutational process can have a qualitative impact on the long-term behavior of an eco-evolutionary system, in contrast to the classical population genetic assumption that all mutations can be modeled with an effective, homogeneous rate. We demonstrate this in the context of a version of MacArthur’s consumer-resource model in which consumers mutate along a resource preference trait-space. Starting from a stochastic individual-based model, we simulate the system in the case where mutations are exogenously generated at a fixed rate (e.g. via external mutagens) and in the case where mutations are coupled to replication (e.g. via DNA copying errors). We find that, surprisingly, replication-coupled mutations are capable of generating a patterned phase in the limit of fast ecological relaxation – precisely the regime where classical population genetic models are expected to operate. We derive the appropriate mean-field description of the stochastic model and use it to show that the patterned phase comes about due to a Turing-like mechanism driven by the non-reciprocal and nonlinear nature of replicative mutations. We demonstrate that these results are robust to demographic noise and model choices and we discuss systems in which this phenomenology might be relevant.

## 1 Introduction

Canonical theories hold that ecology and evolution act on disparate timescales [1], so that ecological and evolutionary processes are often treated as effectively decoupled from each other. However, there is mounting evidence to suggest that ecology and evolution can act on comparable timescales in both natural and laboratory populations, so that these processes couple, generating ‘eco-evolutionary feedback.’ To be more specific, ecological conditions (abiotic factors, interspecies interactions, etc) dictate the course of future evolution. For instance, the presence of particular mutagens might impact the spectrum of mutational signatures that are observed in tumors [2]. Likewise, small evolutionary changes can drive the state of the ecosystem. For example, a monomorphic population can split into two coexisting lineages after the emergence of a few linked single-nucleotide changes [3].

Despite the clear evidence that eco-evolutionary feedbacks can be physically relevant, there remains a dearth of theoretical approaches to describe the phenomenology of eco-evolutionary feedback, in part because eco-evolutionary models must simultaneously account for population growth, nonlinear interactions, mutation, and stochasticity. Many past works that explicitly couple evolution and ecology assume a separation of timescales between the arrival of new phenotypic mutations and the typical equilibration time of the ecological interactions (e.g., the ‘adaptive dynamics’ framework [4], among others [5]). This ‘quasi-static’ approach greatly simplifies analysis, as the dynamics are primarily driven by an invading strain’s linearized growth rate. However, even in this simplified scenario, complex abundance trajectories readily emerge in sufficiently high-dimensional phenotypic spaces [6], raising questions about the general self-consistency of the separation of timescales assumption.

More recently, models of rapid eco-evolutionary dynamics have been proposed, primarily in the context of host-pathogen co-evolution. These have been shown to demonstrate a plethora of interesting behaviors that encompasses and extends the results of adaptive dynamics, without resorting to restrictive assumptions on relative rates. With these models, it has been shown that rapid eco-evolutionary dynamics can generate rich dynamical behavior. For instance, it has been suggested that traveling waves and diversification [7, 8] can be generated by rapid eco-evolutionary processes and that these phenomena might apply to many different contexts. Concurrent work has shown that rapid evolutionary dynamics can buffer against both stochastic [9] and chaotic [10] abundance fluctuations that would otherwise rapidly drive populations extinct.

It has been recently demonstrated that, in the context of host-pathogen co-evolution, distinct species clusters can emerge from otherwise translation-invariant population dynamics on a continuous phenotype space – a phenomenon that has been termed ‘trait-space patterning’ [11]. In Ref. [11], a mutation-independent pattern-formation mechanism was identified that crucially depends on asymmetric, finite-ranged interactions between the pathogen and the host immune system. Trait-space patterning generated by this mechanism can therefore be described as an ecological phenomenon rather than a distinctly eco-*evolutionary* phenomenon, cf Ref. [12]. In the present work, we show that trait space patterning can also emerge in an eco-evolutionary system in a way that explicitly depends on the mutational process.

In particular, we focus on two classes of mutational processes: 1) *exogenous* mutations (i.e. those generated by external mutagens like UV radiation) 2) *replicative* mutations (i.e. those generated by errors intrinsic to the DNA replication process). In particular, exogenous mutations are defined by a rate or *mutation per individual per time*, while replicative mutations are defined by a *probability per birth*. When coupled to the particular coarse-grained constant population size models commonly used in population genetics (e.g. the Wright-Fisher model or the Moran model), these distinct mutational models lead to the same limiting ‘clocklike’ behavior on the timescale of generations, albeit with slightly different relationships between the underlying ‘microscopic’ rate parameters and the ‘macroscopic’ clock rate [13, 14]. As a result, it has oft been assumed that there are no appreciable qualitative differences between these mutational classes, though there is clearly some ambiguity (as pointed out succinctly in Ref. [15]). As we go on to show, in eco-evolutionary contexts, when the supply of mutations is sufficiently large, these two classes of mutational process can drive qualitatively different population-scale behavior.

In Figure 1, we illustrate the simplest scenario in which these two mutational processes can lead to very different outcomes on short timescales. Consider an ecosystem of two ecotypes (or distinguishable subpopulations) and consider that negligible death occurs over our observation window (e.g. *t*_obs_ ≪ typical lifetime of an individual of either subtype). Ecotype A is large and static (i.e. there is no growth or death of A over the observation window) while ecotype B is small and growing so that the total population size *N*_tot_ = *N*_*A*_ + *N*_*B*_ remains approximately constant in log space. In the situation where we only have exogenous mutations, almost all mutations will occur on the static, but large ecotype A background. In contrast, with only replicative mutations, all mutations will occur on the growing but small ecotype B background. These two scenarios, in turn, can lead to very different outcomes on longer timescales, especially if the mutations are beneficial or ecologically relevant. When an eco-evolutionary system consists of many more than two ecotypes and when there are explicit interactions between them, these differences can be amplified even further, particularly in cases where the ecotype abundances follow chaotic trajectories (as can be the case in highly diverse ecosystems).

**Figure 1.**
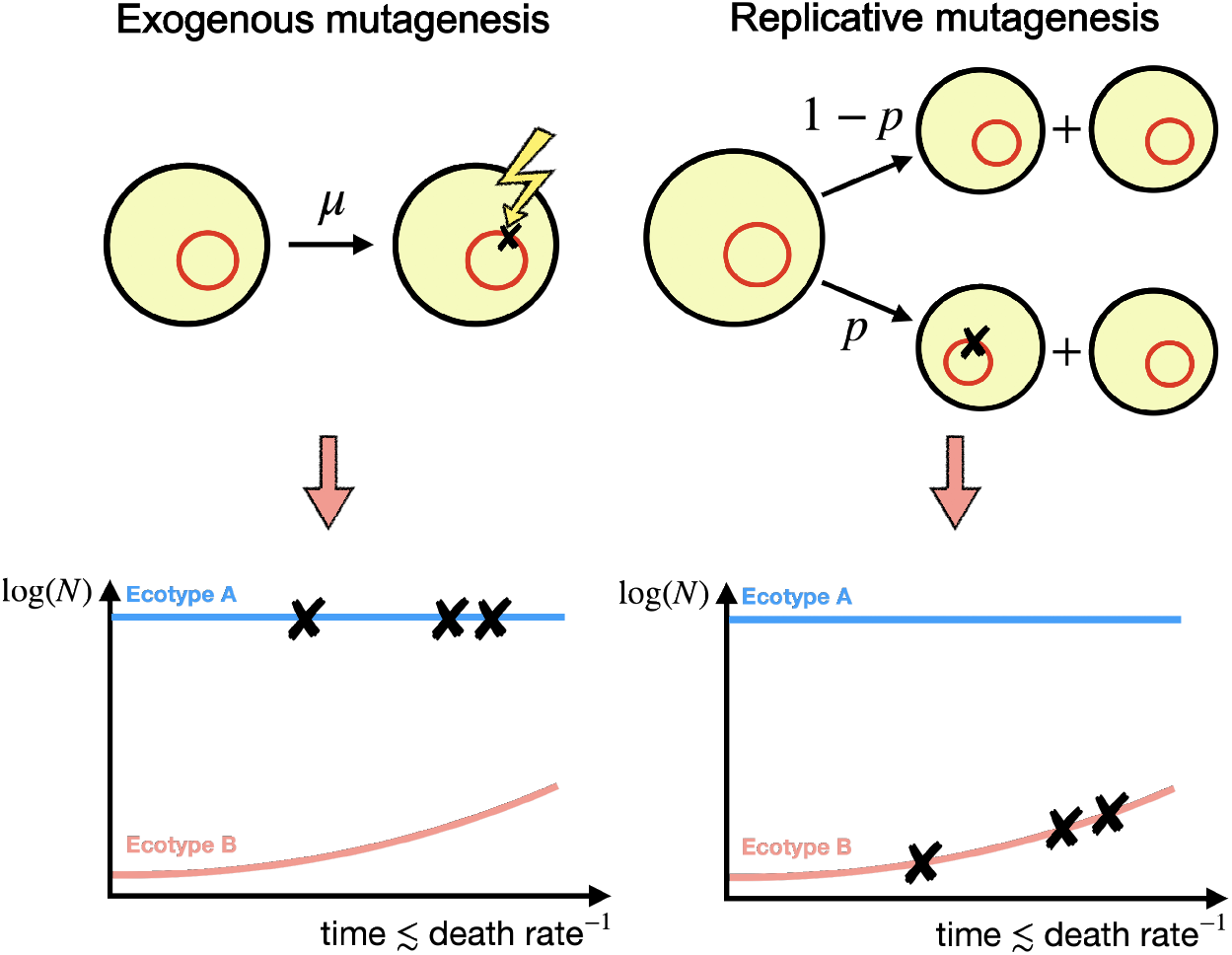
Exogenous and replicative mutations can lead to different outcomes for the same underlying population dynamics. Exogenous mutations preferentially occur on highly abundant ecotype backgrounds, while replicative mutations preferentially occur on growing ecotype backgrounds.

In the remainder of this text, we go on to demonstrate that the details of the mutational process can matter for the long-term behavior of an eco-evolutionary system in the context of MacArthur’s consumer-resource model [16] where we hold the resources fixed and let the consumers evolve rapidly. It stands to be emphasized that the particular consumer-resource model we study (without mutation) is an archetypal example of a *stable* ecology with a homogeneous steady state, so that we do not expect large or chaotic fluctuations due to ecology alone. We show that when mutations are replicative, trait-space patterning emerges in regimes where ecological relaxation and mutation are fast, while no such phase exists for exogenous mutations. Individual-based simulations demonstrating replicative patterning are shown in Fig. 2. We derive a continuum mean-field model from this individual-based description, demonstrating that it captires key elements of the patterned regime of the stochastic dynamics. We discuss the qualitative ingredients required for mutation-induced trait-space patterning due to eco-evolutionary feedback.

**Figure 2.**
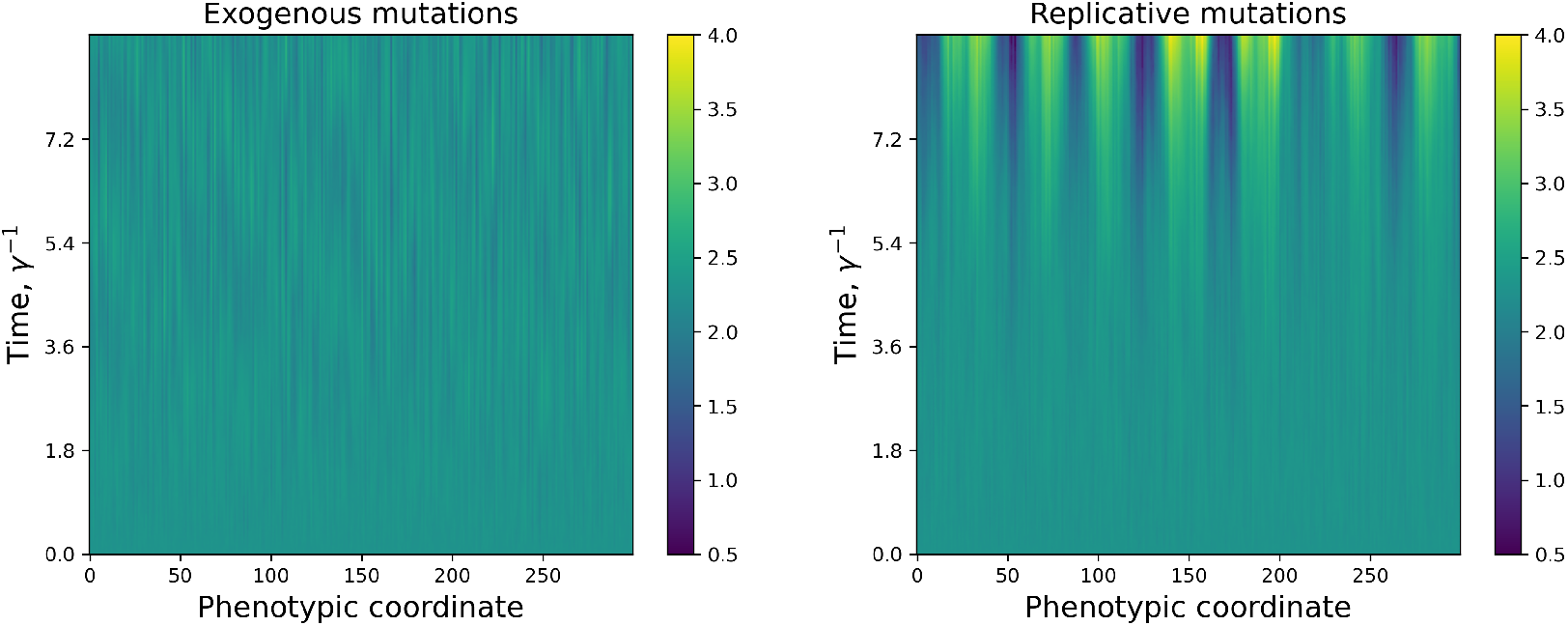
Kymographs of log_10_ consumer abundance from stochastic simulations of the evolving consumer-resource model for exogenous mutations (left panel) and for replicative mutations (right panel). Time is measured in units of the consumer death rate and phenotype space is measured in units of the lattice constant. Mutational jump kernel is the symmetric delta function kernel, 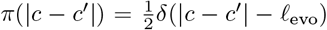, 𝓁_evo_ = 20 in units of the lattice spacing. Both simulations are for a comparable parameter set in the replicative patterned phase (lattice values, *g* = 1*e*6, *κ* = 1*e*7, *γ* = 5*e*5, 𝓁_*Ψ*_ = 𝓁_*ϕ*_ = 1, *Z*_1_ = 5*e*3, *Z*_2_ = 1; exogenous, *μ* = 5*e*5, and replicative, *p* = 1, so that *β*_mut_ = 1/2 for both simulations), initialized to the same uniform initial condition, and run for ~ 9 generations (estimating the generation time as the typical consumer lifetime, 1/*γ*).

## 2 Phenotypic ‘length scales’ of ecology and evolution

In introducing our model, it is important from a conceptual standpoint that we define the space in which it operates and the relevant length scales that define that space. In the present work, we assume that phenotype space is a *d*-dimensional square lattice with lattice spacing *h*, indexed by coordinate *c*. Both consumer and resource types occupy sites on this lattice and interact with each other in a way that depends on their separation. This sort of Euclidean ‘shape space’ model has been invoked to describe the effective spatial extent of all possible host-pathogen pairs [17, 8], where different dimensions might correspond to different antigenic ‘footprints.’ However, this lattice topology is extremely simple, and unlikely to reflect the topology of real phenotypic spaces, which might include trade-offs [18, 19] and intrinsic fitness differences [20]. We nonetheless choose to work with this simple topology to highlight the effect of mutational processes in a way that is isolated from other details of the ecology.

The lattice spacing *h*, intuitively speaking, can be thought of as the smallest relevant unit of change in an eco-evolutionary system. We assume that resource self-limitation occurs on this length scale so that the resources grow according to logistic dynamics at each site. The interpretation is that individual units of resource are exchangeable *with respect to each other* at the length scale of the lattice spacing. We make this choice primarily for convenience in the current exposition, though we are free to set a resource mixing scale based on the requirements of the specific physical system our toy model is to represent. All other length scales are assumed to be ≫ *h*, so that none of our results depend on *h*.

There are then two important long length scales that inform the dynamics. The first is the characteristic length scale on which ecological processes occur, 𝓁_eco_. In our context, this is the typical separation on which consumer preference for resources decays. In other words, this is the length scale on which resources are roughly exchangeable *with respect to the consumers*. For simplicity, we assume that consumer preferences are encoded by a normalized, translationally invariant kernel *Ψ*(|*c* − *c*′|), where *c* is the consumer coordinate and *c*′ is the resource coordinate. We assume that *Ψ* has a characteristic length scale, 𝓁_eco_, in units of the lattice spacing. Many results will hold for different functional forms with the same typical length scale. For much of this work, we will consider interaction kernels with an exponential functional form:

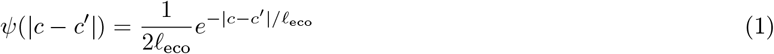

though many results will hold for different functional forms with the same typical length scale.

The second important length scale is the typical length scale on which evolution occurs, 𝓁_evo_. This can be interpreted as the typical distance a mutational event will carry a mutant in phenotype space. We assume that mutational jumps are also encoded by a normalized, translationally invariant kernel, *π*(|*c* − *c*′|), where *c* is the source site and *c*′ is the target site. Many of our results are robust to choices of jump kernel, so long as *π*(|*c* − *c*′|) is finite-ranged. Infinite-ranged *π*(|*c* − *c*′|) will destroy any order in the system, even if *π*(|*c* − *c*′|) has finite moments [21].

## 3 The evolving consumer-resource model

We write down a set of reactions involving the abundances of two types, resources (*R*) and consumers (*N*), which both exist along the discrete, *d*-dimensional phenotype space described in the previous section. The resources grow with rate *g* and saturate at local carrying capacity *κ* in the absence of consumers. As described in the previous section, for the present exposition, we assume that resources do not directly interact across trait-space sites. Consumers die at fixed rate *γ* in the absence of resources.

The consumers and resources are coupled by a consumption process in which a consumer eats the resource on and surrounding its phenotypic coordinate. Consumption is decomposed into two subprocesses, whereby one subprocess consists of events that generate new consumers, while the other subprocess leaves the consumer population size unchanged. The first subprocess is defined by a normalized kernel *Ψ*(|*c* − *c*′|) and a weight *Z*_1_, while the second is defined by a normalized kernel *ϕ*(|*c* − *c*′|) and a weight *Z*_2_. For both kernels, the consumer coordinate is denoted *c* and the resource coordinate is denoted *c*′. Each of these kernels can, in general, decay with phenotypic distance at different rates, or can be of different ranges. As noted earlier, it has been shown that incommensurate, finite-ranged kernels lead to patterning independent of the mutational process [11]. We primarily consider the exponential class of interaction kernels to differentiate our patterning mechanism.

We allow the consumers to mutate according to an exogenous process or a replicative process. For exogenous mutations, mutations occur randomly according to a fixed rate *μ* per individual consumer. Once an individual at a source site is picked, it then jumps to a new phenotypic site with some probability that depends on the distance between the source *c* and target *c*′, *π*(|*c* − *c*′|). For replicative mutations, every birth results in a mutation with some fixed probability *p*. Once a birth event is picked to mutate, the offspring is transported according to the jump kernel, *π*(|*c* − *c*′|). A schematic of the translation invariant model is shown in Figure 3 and the reactions are summarized in Table 1.

**Table 1:** Set of reactions in the model and their resulting continuum descriptions. Terms annotated with (i) appear in the mean field equations as integrals over the coordinate with primes.

| Process | Reaction | Propensity | Continuum |
| --- | --- | --- | --- |
| Resource growth | $R(c) \rightarrow 2R(c)$ | $gR(c)$ | $g\rho(c)$ |
| Resource decay | $2R(c) \rightarrow R(c)$ | $gR^2(c)/\kappa$ | $g\rho^2(c)/\kappa$ |
| Consumption | $N(c') + R(c) \rightarrow N(c')$ | $Z_2\phi(c - c')N(c')R(c)$ | $Z_2\rho(c)\phi(c - c')\chi(c')$ (i) |
| Consumption+birth | $N(c) + R(c') \rightarrow 2N(c)$ | $Z_1\psi(c - c')N(c)R(c')$ | $Z_1\chi(c)\psi(c - c')\rho(c')$ (i) |
| Consumer death | $N(c) \rightarrow \emptyset$ | $\gamma N(c)$ | $\gamma\chi(c)$ |
| Exogenous mutation | $N(c) \rightarrow N(c')$ | $\mu N(c)$ | $\frac{\mu}{2d}\mathcal{S}_\pi\chi(c)$ |
| Replicative mutation | $N(c) \rightarrow N(c) + N(c')$ | $pZ_1\pi(c - c')\psi(c - c'')N(c)R(c'')$ | $\frac{p\gamma}{2d}\frac{1}{\rho^*}\mathcal{S}_\pi\psi(c - c'')\rho(c'')\chi(c)$ (i) |

**Figure 3.**
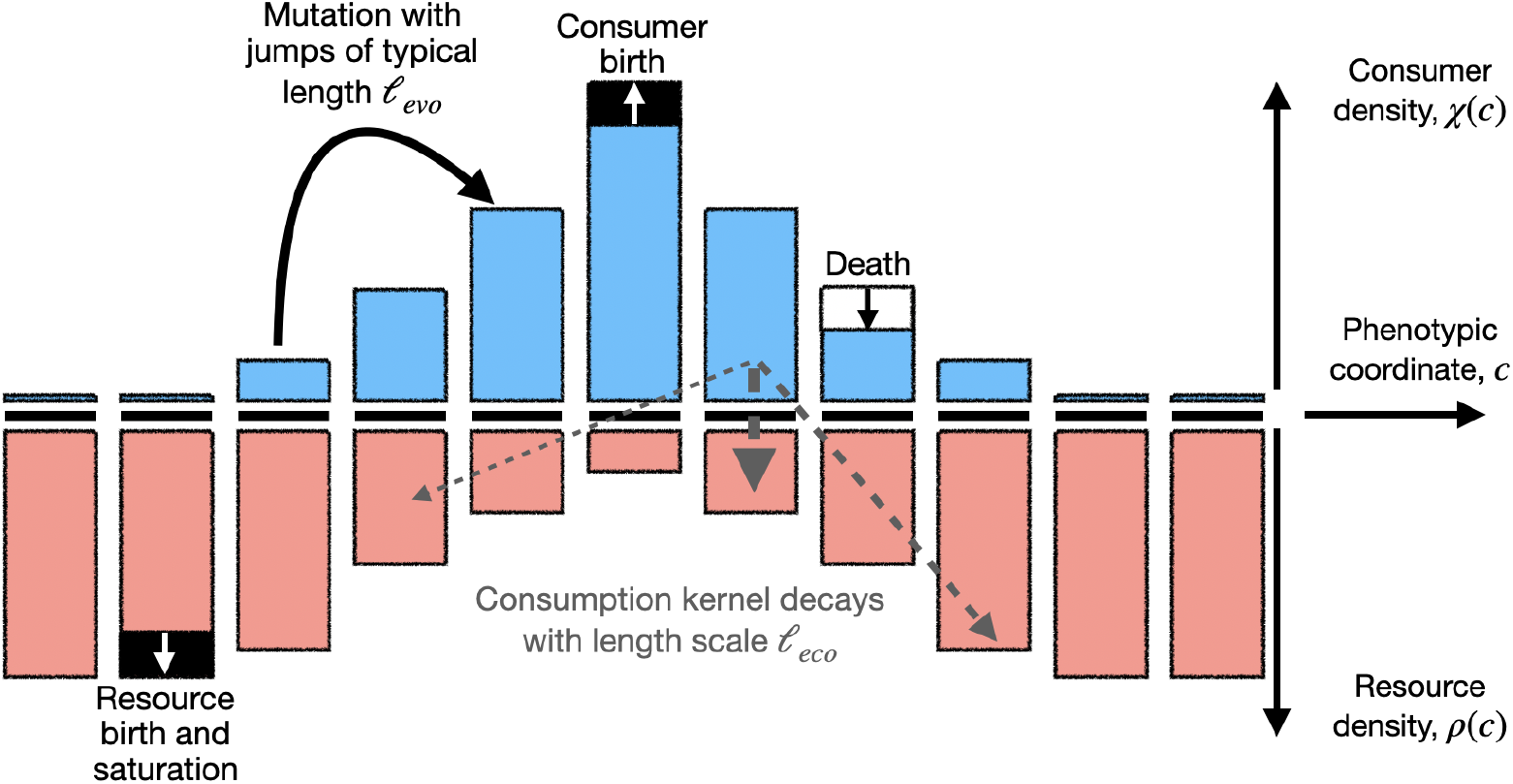
Evolving consumer-resource model schematic, where the height of the bars denotes the density of individuals in an infinitesimal volume of a one-dimensional phenotype space. Consumer individuals die at a fixed rate, grow by consuming resources accourding to a kernel *Z*_1_*Ψ*, consume unproductively according to kernel *Z*_2_*ϕ* and mutate due to external mutagens (‘exogenous mutations’) or DNA replication errors (‘replicative mutations’). Resources grow, compete within a phenotypic volume, and are consumed.

Given that reaction-diffusion processes are difficult to analyze directly at the individual-based level, we leverage the Doi-Peliti framework to construct a path integral (i.e. ‘mesoscopic’) representation of the underlying individual-based process (see [21]). This mathematical object can in turn be used to compute statistics of consumer (*χ*) and resource (*ρ*) concentration fields (abundance per volume of phenotype space). Importantly, this framework allows us to derive a coarse-grained description that is consistent with an underlying individual-based dynamics, rather than relying on ad hoc constructions which may or may not be readily mapped to an underlying microscopic model. The dictionary between underlying reactions and the relevant terms in the continuum model are described in Table 1.

After performing this procedure, we obtain the following mean-field equations for the consumer and resource population densities in phenotype space:

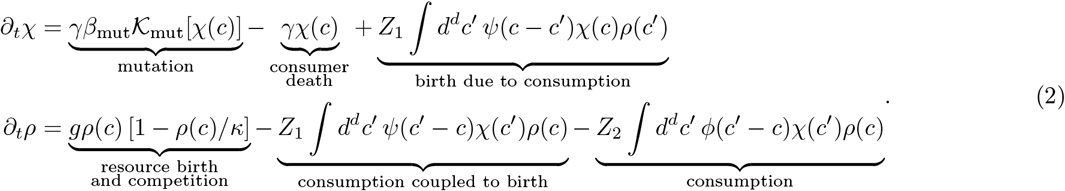

We define *β*_mut_ as the rescaled mutation rate and where 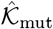 is a normalized mutation kernel, which depends on the class of mutational process and the particular jump kernel we consider. We have the following for exogenous mutagenesis:

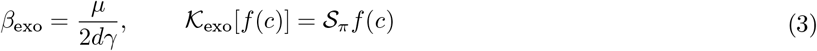

where 𝒮_*π*_ is a differential operator related to the mutational jump process. Whereas, for replicative mutagenesis, we have the following:

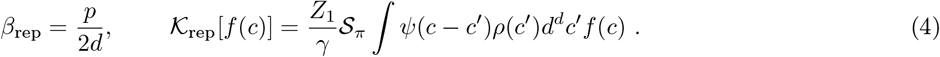

The factors of 2*d* are due to the dimension of the underlying Euclidean lattice. We have that the replicative mutational supply is bounded, *β*_rep_ ≤ 1/2*d*.

We also define the operator 𝒮_*π*_, which encodes the continuum transport properties of the mutational jump kernel *π*. In particular, *S*_*π*_ is related to the Kramers-Moyal expansion [22]:

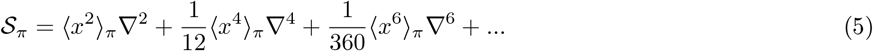

We keep all orders of the expansion because we are observing the system on lengthscales shorter than 𝓁_evo_, the jump length scale. The Fourier transform of 𝒮_*π*_ is related to the characteristic function of the jump kernel 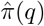:

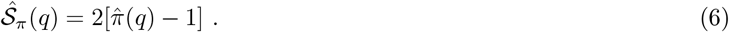

This operator is derived and described in more detail in the SM [21].

## 4 Nondimensionalization

To start, we analyze the mean-field equations assuming identical interaction kernels, 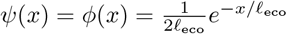. We can write down expressions for the homogeneous fixed point:

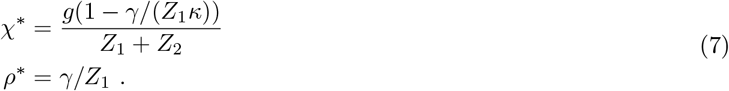

The non-trivial homogeneous fixed point is only valid for *γ/Z*_1_*κ <* 1, where *γ/Z*_1_*κ* = 1 corresponds to a continuous transition. The limit *κ* → ∞ is a singular one in which a local conserved quantity emerges [23] and the ecological dynamics exhibit neutral limit cycle oscillations. For finite *κ*, we will call the quantity *α* ≡ 1 −*γ/Z*_1_*κ* the dimensionless *scaled relaxation rate*. Intuitively, *α* measures the net growth of the consumer population over the timescale that a consumer is typically born at the resource capacity *κ*. This parameter is a measure of the distance from the mean field consumer extinction transition which occurs at *α* = 0 and is inversely proportional to the ecological relaxation timescale *T*_eco_ ∝ *α*^−1^, which diverges near the critical point. Conditioned on *α* > 0, the homogeneous fixed point is stable for all choices of parameters for positive definite kernels *Ψ* and *ϕ* in the absence of mutation. As noted in [11], non-positive definite *Ψ* and *ϕ* can drive patterning in a mutation independent way.

We nondimensionalize Eqs. 2, measuring consumer and resource densities in terms of their fixed point values (*ρ* → *ρ*\**ρ* and *χ* → *χ*\**χ*), measuring time in units of *γ*^−1^, measuring phenotype space in units where the ecological lengthscale 𝓁_eco_ = 1. We define the auxiliary dimensionless quantity *ν* ≡ *g/Z*_1_ and the normalized auxiliary function 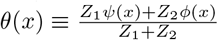 to obtain the following:

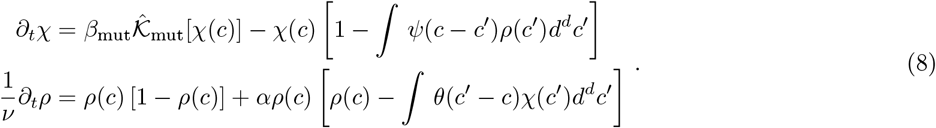

Note, when *ν* ≫ 1, the resource dynamics equilibrate quickly over the timescale *γ*^−1^. Assuming equilibrated resources, the consumer dynamics without mutation obtain a Lyapunov function [24].

## 5 Replicative pattern formation

We now expand the mean field equations around the rescaled fixed point (*χ*\*, *ρ*\*) = (1, 1) and Fourier transform, obtaining the Fourier space Jacobian, **J**(**q**), for both replicative and exogenous mutational kernels, as well as for general mutational kernels, which we report in the SM [21]. Linear stability holds for a particular wavevector **q** if Tr(**J**(**q**)) < 0 and det(**J**(**q**)) > 0 for a two-species system. The trace condition is met for both mutational classes. For exogenous mutagenesis, the Jacobian determinant is the following:

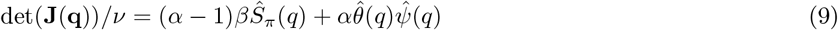

where 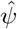 and 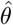 are the Fourier transforms of *Ψ* and *θ*, respectively. Since 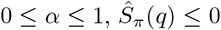 and *Ψ* and *θ* are positive definite by assumption, this expression is always positive. For replicative mutagenesis, the Jacobian determinant is the following:

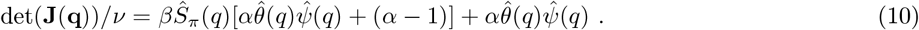

When consumer-resource dynamics are fast (*α* ∼ 1) this inequality simplifies. Given positive definite *θ* and *Ψ*, replicative mutagenesis can achieve det(**J**(**q**)) < 0 when:

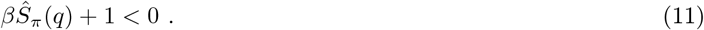

In Fig. 4, we test the prediction for instability using stochastic simulations of the model in Table 1. We observe a patterned phase for sufficiently large *β* with peaks in the consumer and resource abundances out of phase with each other. The instability in Eq. 11 happens for a range of parameter values in the model. Fig. 5 shows the phase diagram generated by inequality 11 for jump kernels of the form 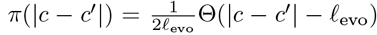 in arbitrary dimensions. Instability can only occur in a region where *β* = *p/*2*d* ≳ 0.43, restricting patterning to one-dimensional phenotype space. In the next section, we discuss modifications to the model that can extend this range, thereby extending patterning to higher dimensions. In the unstable regime, we can use the instability condition to approximate the smallest unstable wavevector:

**Figure 4.**
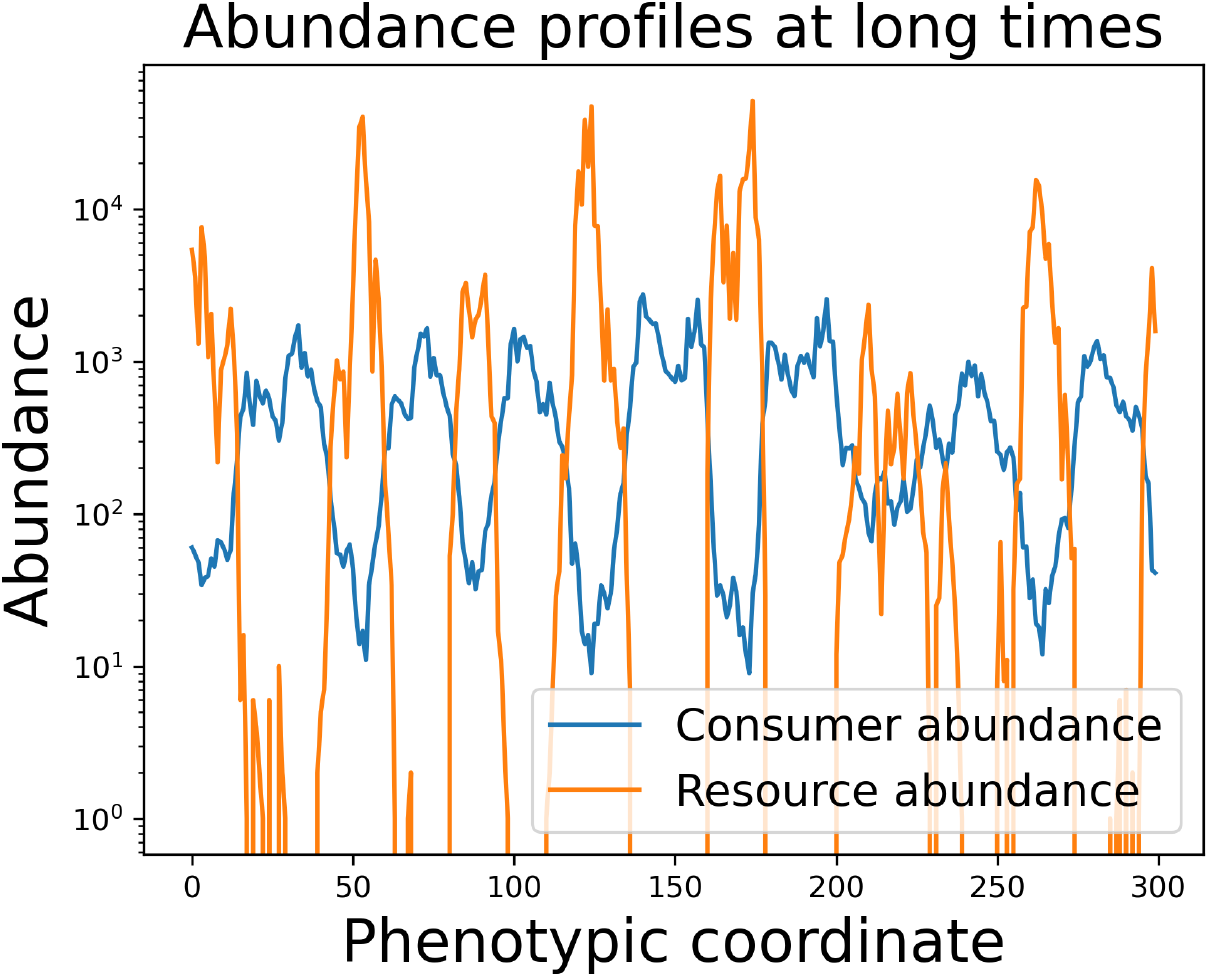
Longtime (*t* ∼ 9*γ*^−1^ units) profile of consumer and resource abundance distributions. Consumer and resource abundance profiles are out of phase with each other. Mutational jump kernel is the uniform step kernel, 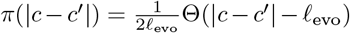, and 𝓁_evo_ = 20 in units of the lattice spacing. Parameters *g* = 1*e*6, *κ* = 1*e*7, *γ* = 5*e*5, 𝓁_*Ψ*_ = 𝓁_*ϕ*_ = 1, *Z*_1_ = 5*e*3, *Z*_2_ = 1, *p* = 1)

**Figure 5.**
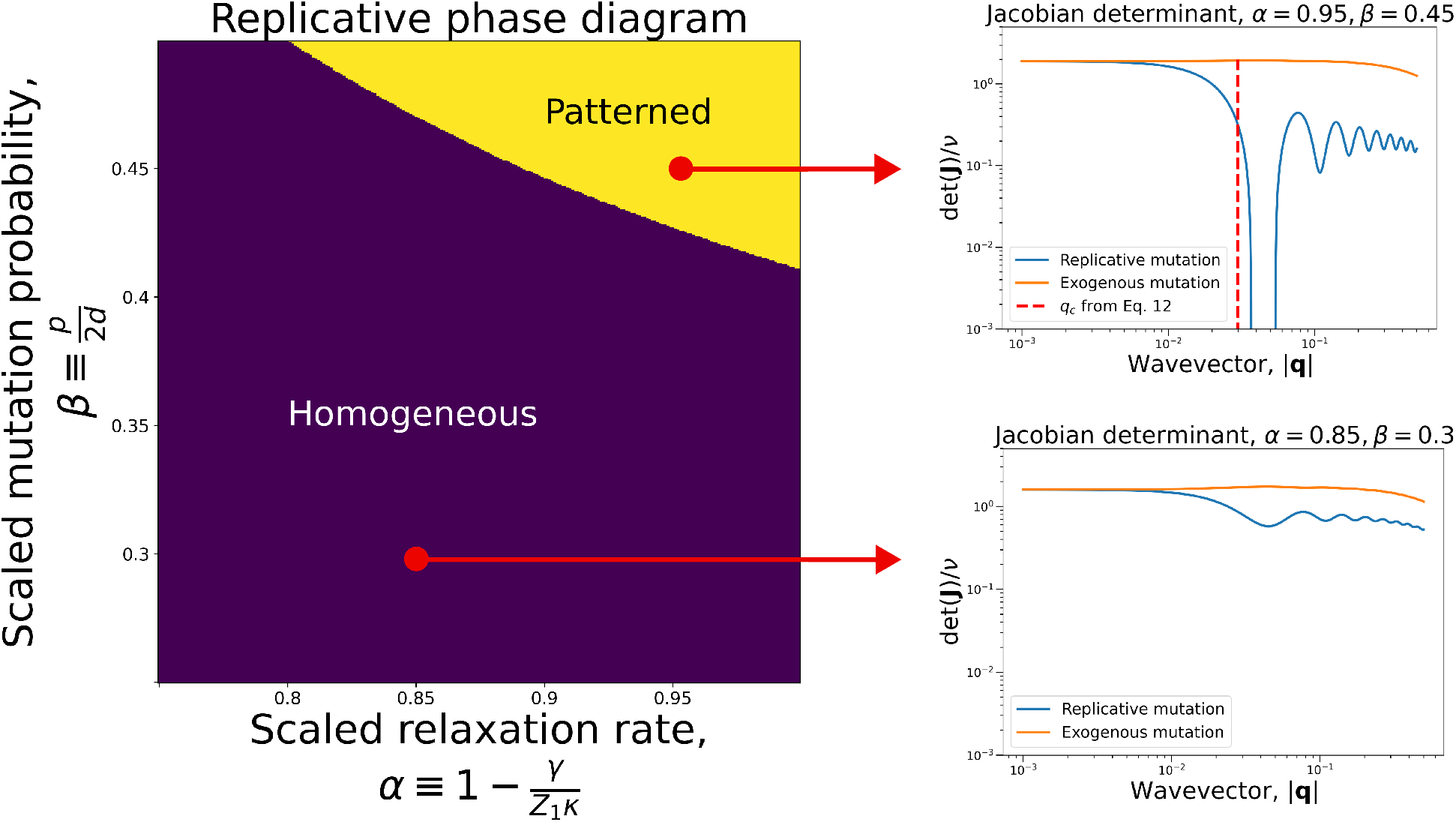
Phase diagram of replicative mean-field pattern formation in the evolving consumer-resource model with step function mutation kernel, 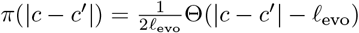, and exponential interaction kernels (Eq. 1). Exogenous mutations do not yield patterns, as evidenced by the positivity of det**J** throughout the phase diagram (orange curves in the right insets). Replicative mutations, on the other hand, can generate patterns for sufficiently high mutation rates and fast relaxation rates (blue curves in the right insets). Note, the right plots are plotted on a log-log scale to emphasize the points where the plotted functions are negative. Individual-based simulations exhibit patterning in the same regions of parameter space predicted by the mean-field model (not shown).

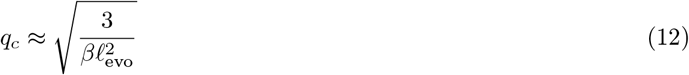

which only depends on the mutation rate and 𝓁_evo_. It should be noted that neither the onset of patterning, nor the lowest unstable wavelength depend on the lengthscale of the ecological interactions 𝓁_eco_.

We say that replicative patterning is Turing-like because it achieves det(**J**(**q**)) < 0. Notably, our mechanism differs from the classical Turing mechanism since only one of the species ‘diffuses’ in our model. However, since its diffusivity is a function of the other species’ abundance, an off-diagonal term is introduced in the Jacobian, leading to patterning. In [21], we show that replicative mutagenesis can yield det(**J**) < 0 for other commonly studied ecological models as well as for certain parameter regimes of models in which the resource can also mutate. In particular, we show that when consumer and resource mutation supply is comparable, patterning is lost and the homogeneous state is achieved, recapitulating observations in Refs. [10, 11].

## 6 General replicative patterning

Let us assume that we have an ecosystem with two classes, *χ* and *ρ*, where *χ* evolves through replicative mutations. We obtain the following mean-field dynamics:

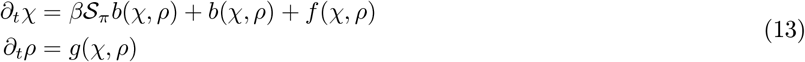

where *b*(*χ, ρ*) is a function describing the birth process of *χ, f* (*χ, ρ*) describes limitation of *χ*, and *g*(*χ, ρ*) describes both birth and limitation of *ρ*. Assume that without mutation (*β* = 0) there is a stable fixed point, (*χ*\*, *ρ*\*).

We define the following matrices, which depend on Fourier transformed derivatives of *b, f*, and *g* and in general depend on *q*:

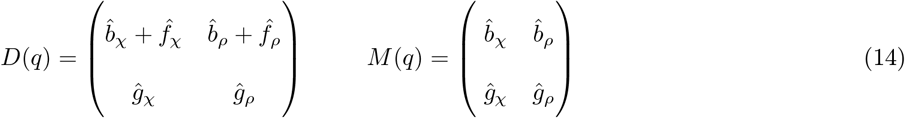

where the matrix *D* is the stability matrix without mutation. The condition for Turing-like instability is then given in terms of the determinants of these matrices given by the inequality:

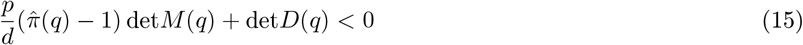

where 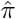 is the characteristic function of the mutational jump kernel *π*. By assumption, the dynamics are stable without mutation, we must have det*D*(*q*) > 0. We also have that 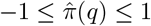 so that the coefficient of det*M* (*q*) is never positive. Therefore, we can only get an instability if det*M* (*q*) is positive and is sufficiently large compared to det*D*(*q*). Given these constraints, we see that if the consumer limitation function *f* depends on both *ρ* and *χ* and its derivatives are negative, we can extend the domain of trait space patterning. For instance, this could be achieved if some resources were ‘poisonous’ to a particular consumer type (e.g. *f* (*χ, ρ*) ∝ − *χ*^*m*^*ρ*^*n*^) – in the context of bacteria and phage such a term might correspond to the presence of defense systems. Describing the details of particular systems that might have larger patterned domains will be explored in future work.

## 7 Discussion

We have described a mechanism by which trait-space patterning might emerge due to mutational processes. In short, the necessary conditions for patterning in the consumer-resource context are as follows: consumer mutation is correlated with replication, it must be long-ranged (𝓁_evo_ ≫ *h*), and it must be frequent (*p* ∼ 1). In addition, the consumer birth process should be sufficiently fast when compared to the death process. This pattern-formation is Turing-like except that it is mediated by a single diffusing field. This field is coupled to a non-diffusing or weakly diffusing resource field via a non-reciprocal diffusivity. We have sketched the phase diagram of the model for particular choices of mutational jump kernels, demonstrating that patterning occurs in a finite region of parameter space. Simulations show that the patterned phase is robust to demographic noise, as is the case in spatially extended ecological models that exhibit pattern formation [25, 26].

It should be noted that the asymmetry of consumption and consumer birth makes the underlying ecological dynamics of our model non-reciprocal in the general case. However, the interactions are still translationally invariant in phenotype space, which facilitates much of our analysis but avoids some of the particular qualitative behaviors that are associated with asymmetry, like chaos [27, 28]. Such chaos-generating asymmetric interactions are not necessary for the patterning phenomenology we describe, though it is unclear how they might deform the results presented here. The case of disordered non-reciprocal interactions was studied in Refs. [5] and [10], primarily with respect to questions about the reachability of the homogeneous phase, and is touched on briefly in the SM [21]. It should also be noted that the relevant non-reciprocal coupling in the present model is generated by nonlinear diffusion due to replicative mutations. This term is conservative, analogous to terms of the non-reciprocal Cahn-Hilliard model described in Ref. [29]. Studying asymmetric and other extensions of our model, including ones with explicit constitutive fitness differences [20], and placing this work in context of other non-reciprocally coupled models present interesting directions for future work. Moreover, it would be interesting to study whether patterning is more permissive in more complex ecologies [30].

Our results are primarily a ‘proof of principle’ that mutational processes can qualitatively impact eco-evolutionary dynamics. However, there are some real world contexts where this phenomenology could be applicable. For instance, it has been observed that *Prochlorococcus*, the most abundant species of photosynthetic ocean-dwelling microbe, exhibits long-lived, discrete microdiversity [31], whereby hundreds of discrete, strongly diverged genomic backbones coexist within a single mixing volume of ocean water. Given their divergence, it can be inferred that these ‘ecotype’ lineages have persisted for many years in the face of fast turnover times and relatively weak mutation rates, which is inconsistent with traditional population genetic models of linear adaptation. However, *Prochlorococcus* are known to be particularly susceptible to phage, as they lack phage defense systems that are found in other bacterial species [32]. In the course of the roughly two day replication time of *Prochlorococcus*, as many as half of the population in a sampling volume can be killed by phage [33]. Moreover, phage are known to mutate much faster than their bacterial prey [34]. Therefore, pattern-formation by bacteria-phage co-evolution could provide a natural explanation for *Prochlorococcus*’ persistent microdiversity. In order to directly test this hypothesis, it will be necessary to carefully analyze the details of the genetic differences between *Prochlorococcus* ecotypes and how they inform phage susceptibility. After characterizing the structure of phenotype space in this way, it might be possible to probe trait-space patterning through laboratory co-evolution experiments.

In addition, there may be signatures of replicative trait-space patterning in the context of cancer and circulating metabolites. Cancers are likely to operate in the fast relaxation regime since a cancer population is, by definition, rapidly growing (*α* ∼ 1). Moreover, somatic human cells are known to typically generate ≳ 10 point mutations per cell division [35], with much higher rates in cancers. It is known that different cancers are prone to exhibit different mutational signatures, owing to different underlying mutational processes [2]. For instance, exogenous processes like smoking or replicative processes like homologous recombination-based repair errors can leave characteristic signatures in the spectrum of tumor mutations. A recent study highlighted that signatures of exogenous and replicative mutations can be distinguished from sequencing data of tissues known to divide at different rates, even if the mutations are of unknown etiology [36]. By measuring the relative contribution of these signatures, one can identify exogenous or replicative mutation dominant tumors. If then, one is able to sequence the genomes of individual cells, our model would predict that cells from tumors dominated by replicative mutational processes would more readily cluster into discrete subclones while exogenously mutated tumor cells would be more continuously distributed in genotype space. While there is extensive cancer genomic data, much of it is bulk sequencing, which can obscure aspects of the underlying clonal structure of tumors. However, there is a growing effort to sequence large quantities of individual cancer cells [37, 38, 39], suggesting that this hypothesis might be directly tested in the near future.

While discussing this work, we should be clear that we are primarily interested in the *long term* behavior of the systems we consider. However, long term can be quite long in evolutionary contexts, when measured in units of human lifetimes [40], and therefore the timescales required to observe the phenomena we describe could potentially be difficult to access even for microbial or cancer cell line experiments. However, there are several phenomena pertaining to eco-evolutionary dynamics that are inherently transient and nonetheless interesting to study, in particular because they are readily observable on the timescale it takes for a student to obtain a PhD, for instance. An example of such phenomena are traveling waves, which have been discussed in other contexts [8, 7], and have even been experimentally probed in a purely evolutionary context [41]. Forthcoming work will address the effect of mutational processes on transient dynamical phenomena like traveling waves.

## 8 Acknowledgments

Dedicated to the memory of Anton Zilman, who passed away in 2024, and with whom S.M. had discussed some early ideas pertaining to this work. S.M. thanks Zachary Sethna, Ignacio Vazquez-Garcia, Nicole Rusk, Jonathan Yuly, Thierry Mora and Shenshen Wang for valuable feedback. S.M. is grateful for support from the Marie-Josée Kravis Fellowship in Quantitative Biology. T.G. was supported by the Schmidt Science Fellowship. This work was supported in part by the National Science Foundation, through the Center for the Physics of Biological Function (PHY-1734030). This work utilized resources from the High Performance Computing Group at Memorial Sloan Kettering Cancer Center.

