## Supplementary Material for "Trait-space patterning is dictated by the tempo and mode of mutation"

November 22, 2024

#### 1 Simulations

We simulate the underlying individual-based stochastic process using the Gibson-Bruck next reaction method [4], which uses a priority queue and a precomputed dependency graph between reactions to speed up each step of the simulation. This method is primarily useful for situations where there are many reacting species with relatively few propensity updates in between reactions. It can achieve substantial speed improvements over the more commonly used Gillespie method, without sacrificing accuracy as is done in approximate methods like tau leaping. We used the open source Python library pqdict [10], an implementation of the priority queue data structure, as the basis of our simulations.

#### 2 Doi-Peliti formalism

##### 2.1 Particulars of phenotype space and relevant phenotypic lengthscales

We are trying to compare two classes of continuum models residing on the same underlying phenotypic space, so it is important that we pay attention to the discretization of that space, the relevant scales of various processes and the system size for our subsequent analysis. Paying attention to these details will be crucial for potentially connecting these models to real systems in the future, especially because 'phenotype space' is a somewhat abstract concept. It is worth mentioning up front that any reasonable phenotype space is subject to constraints that are different from constraints on real spaces from standard physics models. For instance, in phenotype space, it is reasonable to introduce long-ranged ($\gg h$, the lattice lengthscale) interactions that correspond with interactions between strongly diverged phenotypes, or it is reasonable to introduce large jumps in a short time. This has the potential to be confusing. Moreover, we assume that phenotype space is Euclidean in our exposition, though there is no strict reason to believe this is the case. However, many models of phenotype space are as least 'locally' Euclidean, for instance 'energy budget' models [12] in which phenotypes correspond with resource strategies that reside on a simplex with pure strategies on the vertices. Other models are less Euclidean in character (for instance string matching phenotypes [7]). Therefore, we aim to be as transparent about these details as possible in our exposition.

We start with a model that is defined on a lattice with lattice parameter $h$ which sets our minimum unit scale. Loosely speaking $h$ might correspond to the scale of phenotypic differences due to individual nucleotide substitutions. For simplicity, we assume that resource self-limitation occurs on this scale. Then we have that ecological interactions between consumers and resources occur on a lengthscale $\ell_{eco} \gtrsim h$, whereby individuals separated by a distance $d \ll \ell_{eco}$ in phenotype space interact ecologically, whereas individuals separated by a distance $d \gg \ell_{eco}$ barely interact. Intuitively, we can interpret $\ell_{eco}$ as some sort of taxonomic cutoff, or perhaps in a host-pathogen context, as the scale that sets the size of host clusters with a propensity for infection by similar pathogens. It is nonetheless a fixed, physical scale (i.e. not arbitrary). This is encoded in the interaction kernels that decay with phenotypic separation. We have that mutations also occur on a particular phenotypic lengthscale $\ell_{evo}$, which can be thought of as a typical jump size after a mutational event. Finally, our system extent is then a much larger than any of the other length scales in the system $L \gg \ell_{eco/evo} \gg h$. It will be helpful to keep these scales in mind as we proceed.

#### 2.2 Exogenous mutations: mutations as simple diffusion

We have the following master equation:

$$
\begin{aligned}
\partial_t P(\{r_i\},\{c_i\};t) = \sum_i \Bigg\{ & g\left[(r_i-1)P(r_i-1,c_i;t) - r_i P(r_i,c_i;t)\right] \\
& + \frac{g}{\kappa h^d}\left[(r_i+1)r_i P(r_i+1,c_i;t) - r_i(r_i-1)P(r_i,c_i;t)\right] \\
& + \gamma\left[(c_i+1)P(r_i,c_i+1;t) - c_i P(r_i,c_i;t)\right] \\
& + \sum_j \Big[ Z_2\phi(i-j)\left[(r_j+1)c_i P(r_j+1,c_i;t) - r_j c_i P(r_j,c_i;t)\right] \\
& + Z_1\psi(i-j)\left[(r_j+1)(c_i-1)P(r_j+1,c_i-1;t) - r_j c_i P(r_j,c_i;t)\right] \Big] \Bigg\} \\
& + \sum_{\langle i,j\rangle} \frac{\mu}{2d}\pi(|i-j|)\left[(c_i+1)P(r_i,c_i+1,c_j-1;t) - c_j P(r_i,c_i,c_j;t)\right]
\end{aligned} \tag{1}
$$

where $h$ is the (square) lattice constant or minimum phenotypic distance, that will be scaled out in the continuum limit. The last sum is the mutation term, which sums over all connected sites, denoted by $\langle i,j\rangle$. The mutation rate is normalized by a factor $2d$ so that the total flux out of a particular phenotypic site is $\mu$ when summing over all connected sites for a square lattice. Note that different lattices might have slightly different expressions here, but the fact that the coordination scales with the dimension is independent of lattice choice. We keep this factor for this reason.

We define bosonic ladder operators:

$$
[\hat{c}_i,\hat{c}_j^\dagger] = [\hat{r}_i,\hat{r}_j^\dagger] = \delta_{ij}, \qquad [\hat{c}_i^\dagger,\hat{r}_j^\dagger] = [\hat{c}_i^\dagger,\hat{r}_j] = [\hat{c}_i,\hat{r}_j^\dagger] = [\hat{c}_i,\hat{r}_j] = 0 \tag{2}
$$

We use these to define the state vector:

$$
|\Psi\rangle \equiv \sum P(\{r_i\},\{c_i\};t)\prod_i \hat{c}_i^{\dagger c_i}\hat{r}_i^{\dagger r_i}|0\rangle
$$

which allow us to cast the master equation in a Fock space representation with quasi-Hamiltonian:

$$
\partial_t|\Psi\rangle = -(H_{mut}+H_{int})|\Psi\rangle
$$

which we have decomposed into mutation and interaction terms. We note that we can formally solve this equation to obtain:

$$
|\Psi(t)\rangle = e^{-Ht}|\Psi(0)\rangle
$$

We can write down the interaction Hamiltonian for the above master equation where we have 'normal ordered' operators using the commutation relations so that all daggered variables are to the left:

$$
\begin{aligned}
H_{int}[c,c^\dagger,r,r^\dagger] = & -\sum_i \left[ g(r_i^\dagger-1)r_i^\dagger r_i + \frac{g}{\kappa h^d}(1-r_i^\dagger)r_i^\dagger r_i^2 + \gamma(1-c_i^\dagger)c_i \right] \\
& -\sum_{i,j}\left[ Z_1\psi(i-j)(c_i^\dagger - r_j^\dagger)c_i^\dagger c_i r_j + Z_2\phi(i-j)(1-r_j^\dagger)c_i^\dagger c_i r_j \right]
\end{aligned} \tag{3}
$$

Now we add exogenous mutations, which have the following operator representation:

$$H_{mut} = \frac{\mu}{2d} \sum_{\langle i,j \rangle} \pi(|i-j|)(c_i^\dagger - c_j^\dagger)(c_i - c_j) \ .$$

Taking these together and operating in the space of coherent states, which are eigenstates of the lowering operators, we can replace operators with continuous (complex) eigenvalues:

$$\begin{aligned} c_i &\to h^d \chi_i & \qquad r_i &\to h^d \rho_i \\ c_i^\dagger &\to \chi_i^* & \qquad r_i^\dagger &\to \rho_i^* \end{aligned}$$

where unstarred fields have been rescaled by phenotype space so that now they are densities, while starred noise fields remain dimensionless. We can then follow the standard procedure of first dividing time into discrete chunks, inserting resolutions of the identity at each time step using the over-completeness of coherent states. From here, we take the appropriate continuum limit in time while discarding boundary terms because we are interested in the long time behavior (though these may be important to keep when studying large deviations, for instance). Note that the details of these steps are important, though somewhat cumbersome to describe in detail. We have followed the approaches laid out in much detail in [3, 11, 2] among others.

This procedure gives averages of observables in terms of a path integral:

$$\langle \mathcal{O}(t) \rangle \propto \int \prod_i d\chi_i d\chi_i^* d\rho_i d\rho_i^* \mathcal{O}(\{\chi_i\}, \{\rho_i\}) e^{-S[\chi_i, \chi_i^*, \rho_i, \rho_i^*, t_f]}$$

with action in discrete space given by:

$$S[\chi_i, \chi_i^*, \rho_i, \rho_i^*, t] = \sum_i \int_0^{t_f} dt \ \{\chi^* \partial_t + H_{mut}[\chi, \chi^*, \rho, \rho^*] + H_{int}[\chi, \chi^*, \rho, \rho^*]\}$$

We take the 'naive' continuum limit $\sum_i h^d \to \int d^d x$, $i \to x$. Assuming sufficient smoothness over length scales longer than $h$, expand differences around position $x$, for instance:

$$\chi^\dagger(x') - \chi^\dagger(x) = (x'-x)\nabla\chi^\dagger(x) + \frac{(x'-x)^2}{2!}\nabla^2\chi^\dagger(x) + ...$$

and the same for the observable field $\chi$. This expansion is analogous to performing a Kramers-Moyal expansion of the mutational jump process. Integrating by parts to move differential operators from $\chi^\dagger$ to $\chi$ and using rotational invariance to get rid of odd gradient terms, we get for the continuum mutation term:

$$\chi^\dagger(x) \sum_{\substack{n \equiv m \\ (\text{mod } 2)}}^{\infty} (-1)^{n+1} \int d^d x' \pi(|x-x'|) \frac{|x-x'|^{n+m}}{n!m!} \nabla^{n+m} \chi(x) = \chi^\dagger(x) \sum_{\substack{n \equiv m \\ (\text{mod } 2)}}^{\infty} (-1)^{n+1} \frac{\langle x^{n+m} \rangle_\pi}{n!m!} \nabla^{n+m} \chi(x)$$

where $\langle x^{n+m} \rangle_\pi$ should be interpreted as conditional moments of the jump kernel $\pi$. We define the full continuum jump operator:

$$\mathcal{S}_\pi \equiv \sum_{\substack{n \equiv m \\ (\text{mod } 2)}}^{\infty} (-1)^{n+1} \frac{\langle x^{n+m} \rangle_\pi}{n!m!} \nabla^{n+m}$$

This gives a bulk continuous action in terms of continuous fields:

$$S[\chi, \chi^*, \rho, \rho^*] = \int d^d x \int_0^{t_f} dt \left\{ \chi^* \left( \partial_t - \frac{\mu}{2d} \mathcal{S}_\pi \right) \chi + H_{int}[\chi, \chi^*, \rho, \rho^*] \right\}$$

The phenotype space is now measured in units with $h = 1$, so there is a natural cutoff (which becomes particularly relevant when we examine the system in Fourier space).

To quickly extract the mean field equation from this action, we make a field shift, $\chi^* = 1 + \tilde{\chi}$, $\rho^* = 1 + \tilde{\rho}$, which comes about by commuting operators through the action in the field theory interpretation, or alternatively, perhaps more intuitively, subtracting out the mean from the noise. We get:

$$\begin{aligned} S[\chi, \tilde{\chi}, \rho, \tilde{\rho}] = \int d^d x \int_0^{t_f} dt \Big\{ & \tilde{\chi} \left( \partial_t - \frac{\mu}{2d} \mathcal{S}_\pi \chi(x) + \gamma \right) \chi \\ & + \tilde{\rho} \left( \partial_t - g \right) \rho - g \tilde{\rho}^2 \rho + \frac{g}{\kappa} (1 + \tilde{\rho}) \tilde{\rho} \rho^2 \\ & - Z_1 \int d^d y \psi(x - y) (\tilde{\chi}(x) - \tilde{\rho}(y)) (1 + \tilde{\chi}(x)) \chi(x) \rho(y) \\ & + Z_2 \int d^d y \phi(x - y) \tilde{\rho}(y) (1 + \tilde{\chi}(x)) \chi(x) \rho(y) \Big\} \end{aligned} \tag{4}$$

We can interpret this action as representing a pair of coupled Langevin equations with positive correlators ($\sim \tilde{x_i}^2$), negative cross-correlators ($\sim \tilde{x_i} \tilde{x_j}$) and *without* diffusion noise ($\sim x_i (\nabla \tilde{x_i})^2$). The Langevin equations are given by:

$$\begin{aligned} \partial_t \chi &= \frac{\mu}{2d} \mathcal{S}_\pi \chi(x) - \gamma \chi(x) + Z_1 \int d^d y \, \psi(x - y) \chi(x) \rho(y) + \eta(t) \\ \partial_t \rho &= g \rho(y) \left[ 1 - \rho(y)/\kappa \right] - Z_1 \int d^d x \, \psi(x - y) \chi(x) \rho(y) \\ &\quad - Z_2 \int d^d x \, \phi(x - y) \chi(x) \rho(y) + \xi(t) \end{aligned} \tag{5}$$

with noise correlators:

$$\begin{aligned} \langle \eta(x, t) \eta(x', t') \rangle &= 2 Z_1 \int d^d y \, \psi(x - y) \chi(x) \rho(y) \delta(x - x') \delta(t - t') \\ \langle \xi(x, t) \xi(x', t') \rangle &= 2 g \rho(x) \left[ 1 - \rho(x)/\kappa \right] \delta(x - x') \delta(t - t') \\ \langle \eta(x, t) \xi(y, t') \rangle &= - \Big[ Z_1 \psi(x - y) \chi(x) \rho(y) \\ &\qquad + Z_2 \phi(x - y) \chi(x) \rho(y) \Big] \delta(t - t') \end{aligned} \tag{6}$$

This set of equations is similar to those studied in ecological contexts that involve real space [2].

It is important to note that these Langevin equations are *not the same* as those that are obtained from a system size expansion, and the noise terms are not the right ones to decribe a density field, although the mean field exactly corresponds. In a later section, we will obtain linearized Langevin equations by Cole-Hopf transformation and expansion around the homogeneous steady state of the mean-field model.

#### 2.3 Replicative mutations: mutations as nonlinear diffusion

The derivation will be similar to the one for exogenous mutagenesis, except that we will have to keep track of the fact that a fraction of consumer births result in mutations (so the total number of births is comparable between the replicative and exogenous models). After carefully accounting for terms, we end up with the same interaction term:

$$\begin{aligned} H_{int} = &- \sum_i \left[ g(r_i^\dagger - 1) r_i^\dagger r_i + \frac{g}{\kappa h^d} (1 - r_i^\dagger) r_i^\dagger r_i^2 + \gamma (1 - c_i^\dagger) c_i \right] \\ &- \sum_{i,j} \left[ Z_1 \psi(i - j)(c_i^\dagger - r_j^\dagger) c_i^\dagger c_i r_j + Z_2 \phi(i - j)(1 - r_j^\dagger) c_i^\dagger c_i r_j \right] \end{aligned} \tag{7}$$

and the mutation term:

$$H_{mut} = \frac{pZ_1}{2d} \sum_{\langle i,k \rangle, j} \pi(|k-i|)(c_k^\dagger - c_i^\dagger)[\psi(i-j) c_i^\dagger c_i r_j - \psi(k-j) c_k^\dagger c_k r_j] \tag{8}$$

We take the continuum limit as before and replace discrete operators with continuous fields, expanding discrete differences, to get the following action in units where $h = 1$:

$$\begin{aligned} S[\chi, \chi^*, \rho, \rho^*] = \int d^d x \int_0^{t_f} dt \Big\{ & \tilde{\chi} \partial_t \chi + \gamma(1 - \tilde{\chi})\chi \\ & \tilde{\rho}\partial_t \rho - g(\tilde{\rho} - 1)\tilde{\rho}\rho - \frac{g}{\kappa}(1 - \tilde{\rho})\tilde{\rho}\rho^2 \\ & - Z_1 \int d^d y \, \psi(x-y)(\tilde{\chi}(x) - \tilde{\rho}(y))\tilde{\chi}(x)\chi(x)\rho(y) \\ & + Z_2 \int d^d y \, \phi(x-y)(1 - \tilde{\rho}(y))\tilde{\chi}(x)\chi(x)\rho(y) \\ & - \tilde{\chi}(x) \frac{p\gamma}{2d} \frac{1}{\rho^*} \mathcal{S}_\pi \int d^d y \, \psi(x-y)\rho(y)\tilde{\chi}(x)\chi(x) \end{aligned} \tag{9}$$

where we see the same jump operator $\mathcal{S}_\pi$ which was defined for exogenous mutations.

After Doi shifting, we can derive the coupled 'naive' Langevin equations:

$$\begin{aligned} \partial_t \chi &= \frac{p\gamma}{2d} \frac{1}{\rho^*} \mathcal{S}_\pi \int d^d y \, \psi(x-y)\rho(y)\chi(x) \\ &\quad + Z_1 \int d^d y \, \psi(x-y)\chi(x)\rho(y) + \eta(t) \\ \partial_t \rho &= g\rho(y) \left[1 - \rho(y)/\kappa\right] - Z_1 \int d^d x \, \psi(x-y)\chi(x)\rho(y) \\ &\quad - Z_2 \int d^d x \, \phi(x-y)\chi(x)\rho(y) + \xi(t) \end{aligned} \tag{10}$$

with noise correlators:

$$\begin{aligned} \langle \eta(x,t)\eta(x',t') \rangle &= 2 \left[ \frac{p\gamma}{2d} \frac{1}{\rho^*} \mathcal{S}_\pi \int d^d y \, \psi(x-y)\rho(y)\chi(x) \right. \\ &\qquad \left. + Z_1 \int d^d y \, \psi(x-y)\chi(x)\rho(y) \right] \delta(x-x')\delta(t-t') \\ \langle \xi(x,t)\xi(x',t') \rangle &= 2g\rho(x) \left[1 - \rho(x)/\kappa\right] \delta(x-x')\delta(t-t') \\ \langle \eta(x,t)\xi(y,t') \rangle &= - \left[ Z_1 \psi(x-y)\chi(x)\rho(y) \right. \\ &\qquad \left. + Z_2 \phi(x-y)\chi(x)\rho(y) \right] \delta(t-t') \end{aligned} \tag{11}$$

Note that, as above, although this Langevin equation is physically valid it does not correspond to the fluctuating dynamics of a density field. However, as before, the mean field does correspond with the mean field of the density of the underlying individual based stochastic process (see [3] for a succinct discussion of this subtlety).

##### 2.4 Details of the jump operator

We had previously defined the jump operator:

$$\mathcal{S}_\pi = \sum_{\substack{n \equiv m \\ (\mathrm{mod}\ 2)}}^{\infty} (-1)^{n+1} \frac{\langle x^{n+m} \rangle_\pi}{n!m!} \nabla^{n+m}$$

where the sum starts at $n, m = 1$. Upon rearranging this sum, we obtain a more readily interpretable form:

$$\mathcal{S}_\pi = 2 \sum_{n \geq 2,\ \mathrm{even}} \frac{\langle x^n \rangle_\pi}{n!} \nabla^n$$

Therefore the jump operator $\mathcal{S}_\pi$ is analogous to a Kramers-Moyal expansion and consistes of terms of all orders of derivatives. The Fourier dual of this operator can be expressed in simple terms as:

$$\hat{\mathcal{S}}_\pi(q) = \langle\, 2(\cos(qx) - 1)\, \rangle_{\pi(x)} = 2(\hat{\pi}(q) - 1)$$

where $\hat{\pi}(q)$ is the characteristic function of the jump kernel $\pi$. In general, will simply write $\mathcal{S}_\pi$ for the real space operator and will denote the Fourier dual by $\hat{\mathcal{S}}_\pi(q)$.

#### 3 Mean-field analysis

We start by analyzing the deterministic mean-field behavior of the model. We can solve for the homogeneous fixed point for both replicative and exogenous mutagenesis:

$$\rho^* = \gamma / Z_1 \qquad \chi^* = \frac{g(1 - \gamma/(Z_1\kappa))}{Z_1 + Z_2}$$

Note that there is a valid non-trivial homogeneous fixed point only if the condition $(Z_1\kappa)/\gamma - 1 > 0$ is satsified. There is a saddle node bifurcation at $(Z_1\kappa)/\gamma = 1$, causing the non-trivial global fixed point to disappear. As the system is driven towards this bifurcation, it experiences critical slowing down by way of the scaled relaxation rate:

$$\alpha = \frac{Z_1\kappa - \gamma}{Z_1\kappa}$$

We rescale time by the consumer death rate, and rescale the consumer and resource abundances by their fixed point values to obtain the rescaled mean-field equations reported in the main text:

$$\begin{aligned} \partial_t \chi &= \beta_{mut} \hat{\mathcal{K}}_{mut}[\chi(c)] - \chi(c) \left[1 - \int \psi(c - c')\rho(c')d^d c'\right] \\ \frac{1}{\nu}\partial_t \rho &= \rho(c)\left[1 - \rho(c)\right] + \alpha\rho(c)\left[\rho(c) - \int \theta(c' - c)\chi(c')d^d c'\right] \end{aligned} \tag{12}$$

with $\nu \equiv \frac{g}{Z_1}$ and $\theta(x) \equiv \frac{Z_1\psi(x) + Z_2\phi(x)}{Z_1 + Z_2}$. We will use these rescaled equations to describe the stability properties of the fixed point.

##### 3.1 Exogenous mutagenesis

For exogenous mutagenesis, we have the Jacobian matrix:

$$\mathbf{J} = \begin{pmatrix} \beta\hat{\mathcal{S}}_\pi(q) & \hat{\psi} \\ -\nu\alpha\hat{\theta} & \nu(\alpha - 1) \end{pmatrix} \tag{13}$$

The eigenvalues are given by:

$$\omega(\mathbf{q}) = \mathrm{tr}(\mathbf{J})/2 \pm \sqrt{(\mathrm{tr}(\mathbf{J})/2)^2 - \det(\mathbf{J})}$$

with trace and determinant given by:

$$\begin{aligned}
\mathrm{tr}(\mathbf{J}) &= \beta\hat{\mathcal{S}}_\pi(q) + \nu(\alpha - 1) \\
\det(\mathbf{J}) &= \nu\left[(\alpha-1)\beta\hat{S}_\pi(q) + \alpha\hat{\theta}(q)\hat{\psi}(q)\right] .
\end{aligned}$$

#### 3.2 Replicative mutagenesis

For replicative mutagenesis, we have the Fourier transformed Jacobian matrix:

$$\mathbf{J} = \begin{pmatrix} \beta\hat{\mathcal{S}}_\pi(q) & \hat{\psi}\left(1 + \beta\hat{\mathcal{S}}_\pi(q)\right) \\ -\nu\alpha\hat{\theta} & \nu(\alpha-1) \end{pmatrix} \tag{14}$$

where we have:

$$\begin{aligned}
\mathrm{tr}(\mathbf{J}) &= \beta\hat{\mathcal{S}}_\pi(q) + \nu(\alpha - 1) \\
\det(\mathbf{J}) &= \nu\left[\beta\hat{S}_\pi(q)[\alpha\hat{\theta}(q)\hat{\psi}(q) + (\alpha-1)] + \alpha\hat{\theta}(q)\hat{\psi}(q)\right] .
\end{aligned}$$

#### 3.3 Bifurcations

There are two transitions, corresponding to the eigenvalues gaining an imaginary part (global Hopf bifurcation, $\mathrm{tr}(\mathbf{J}(\mathbf{q}=0)) \to 0$) and another corresponding to linear (Turing-like) instability for a set of $\mathbf{q}$ with finite support ($\det(\mathbf{J}) < 0$).

##### 3.3.1 Hopf bifurcation

We discuss the Hopf bifurcation briefly, as it is a standard characteristic of Lotka-Volterra type equations as the carrying capacity goes to zero. We have in general (assuming normalized kernels WLOG so that $\hat{\psi}(0) = \hat{\phi}(0) = 1$):

$$\omega(\mathbf{q}=0) = -\frac{\nu(\alpha-1)}{2} \pm \sqrt{\left(\frac{\nu(\alpha-1)}{2}\right)^2 - \nu\alpha}$$

so that as $\kappa \to \infty$ (or $\alpha \to 1$), a global damped oscillation is developed which approaches (undamped) oscillation frequency in the limit:

$$\omega(\mathbf{q}=0) = \pm i\sqrt{\nu}$$

This is due to a conserved quantity emerging in the limit, which has been discussed elsewhere.

##### 3.3.2 Turing instability

For exogenous mutagenesis, we have that Turing instability is given in the general case by:

$$(\alpha-1)\beta\hat{S}_\pi(q) + \alpha\hat{\theta}(q)\hat{\psi}(q) < 0 . \tag{15}$$

For replicative mutagenesis, we have the instability condition:

$$\beta\hat{S}_\pi(q)[\alpha\hat{\theta}(q)\hat{\psi}(q) + (\alpha-1)] + \alpha\hat{\theta}(q)\hat{\psi}(q) < 0 . \tag{16}$$

##### 3.3.3 Turing analysis for $\alpha \ll 1$

For the regime $\alpha \ll 1$, the Turing conditions for replicative and exogenous mutagenesis are similar. The following approximate inequality must hold:

$$-\beta \hat{S}_\pi(q) + \alpha \hat{\theta}(q)\hat{\psi}(q) \lesssim 0 \tag{17}$$

We have $\hat{\theta}(q)\hat{\psi}(q) \leq 1$ by normalization of $\psi$ and $\theta$. Similarly $\hat{\mathcal{S}}_\pi(\mathbf{q}) = 2(\hat{\pi} - 1)$ is bounded to the domain $[-4, 0]$ since $\pi$ is normalized. Therefore, the only way to achieve the inequality is if $\hat{\theta}(q)\hat{\psi}(q) < 0$ for some range of $\mathbf{q}$. This effect is the same reported in [6].

##### 3.3.4 Turing analysis for $\alpha \gg 1$

For the regime $\alpha \gg 1$, the Turing conditions for replicative and exogenous mutagenesis differ from each other. For exogenous mutations the Turing instability condition does not meaningfully change – only incommensurate, finite ranged interactions can mediate pattern formation. For replicative mutations, however, we have the approximate condition:

$$\alpha \hat{\theta}(q)\hat{\psi}(q) \left[\beta \hat{S}_\pi(q) + 1\right] < 0 \ . \tag{18}$$

Given positive definite $\theta$ and $\psi$, the condition simplifies to:

$$\beta \hat{S}_\pi(q) + 1 < 0$$

As above, $\hat{\mathcal{S}}_\pi(\mathbf{q}) = 2(\hat{\pi} - 1)$ is bounded to the domain $[-4, 0]$. If $\pi$ is positive definite $\hat{\mathcal{S}}_\pi(\mathbf{q})$ is bounded to the domain $[-2, 0]$. Since $\beta = \frac{p}{2d} \leq \frac{1}{2}$, this inequality can only be met for non-positive definite (i.e. finite-ranged) $\pi$.

### 4 Model variants

Here we will study properties of related eco-evolutionary models that might be of relevance in certain contexts. We will restrict our analysis to the mean-field level for simplicity, leaving a more detailed analysis of stochastic variants to future work.

#### 4.1 Generalized mutational processes

For generalized mutation kernels of the form $\mathcal{D}_{mut}(\rho, \chi)$, we have the mean field equations:

$$\begin{aligned} \partial_t \chi &= \mathcal{S}_\pi \mathcal{D}_{mut}(\rho(c), \chi(c)) - \chi(c)\left[1 - \int \psi(c - c')\rho(c')d^d c'\right] \\ \frac{1}{\nu}\partial_t \rho &= \rho(c)\left[1 - \rho(c)\right] + \alpha\rho(c)\left[\rho(c) - \int \theta(c' - c)\chi(c')d^d c'\right] \end{aligned} \tag{19}$$

we have that the Jacobian is:

$$\mathbf{J} = \begin{pmatrix} \partial_\chi \mathcal{D}_{mut}(\rho^*, \chi^*)\hat{\mathcal{S}}_{\pi,\chi}(q) & \tilde{\psi} + \partial_\rho \mathcal{D}_{mut}(\rho^*, \chi^*)\hat{\mathcal{S}}_{\pi,\chi}(q) \\ -\nu\alpha\hat{\theta} & \nu(\alpha - 1) \end{pmatrix} . \tag{20}$$

We see that an off-diagonal term introduced by $\rho$-dependence of the mutational diffusion constant is crucial for nonlinear patterning (as is the case for replicative mutations).

#### 4.2 Mutating resources

In all generality we can have that the resources also evolve. We assume that resources are transported with jump kernel $\xi$ that we obtain the following dimensionless dynamics:

$$\begin{aligned} \partial_t \chi &= \mathcal{S}_\pi \mathcal{D}_{mut,\chi}(\rho(c), \chi(c)) - \chi(c) \left[1 - \int \psi(c-c')\rho(c')d^dc'\right] \\ \frac{1}{\nu}\partial_t \rho &= \mathcal{S}_\xi \mathcal{D}_{mut,\rho}(\rho(c), \chi(c)) + \rho(c)\left[1-\rho(c)\right] + \alpha\rho(c)\left[\rho(c) - \int \theta(c'-c)\chi(c')d^dc'\right] \end{aligned} \tag{21}$$

with the same homogeneous fixed point as before. However, with resources, as they are set up, there is no ambiguity between the scaling of the mutation term, since resource births are proportional to the population size. Therefore, we have:

$$D_{mut,\rho}\left[\chi(c), \rho(c)\right] = \frac{D^{const}_{mut,\rho}}{2d}\rho(c)$$

where for exogenous mutations $D^{const}_{mut,\rho} = \mu$, as before, and for replicative mutations $D^{const}_{mut,\rho} = pg$ with $0 \leq p \leq 1$. We have the Jacobian:

$$\mathbf{J} = \begin{pmatrix} \partial_\chi \mathcal{D}_{mut}(\rho^*, \chi^*)\hat{\mathcal{S}}_{\pi,\chi}(q) & \tilde{\psi} + \partial_\rho \mathcal{D}_{mut}(\rho^*, \chi^*)\hat{\mathcal{S}}_{\pi,\chi}(q) \\ -\nu\alpha\hat{\theta} & \nu(\alpha-1) \end{pmatrix}. \tag{22}$$

where we see that comparable resource mutation favors the homogeneous state, as observed in other work [6, 7], which stems from Turing's original observation that there must be a timescale separation between activator and inhibitor dispersal for patterns to form.

#### 4.3 Externally supplied resource

We consider a system in which resources are externally supplied and lost at a fixed washout rate. These types of models have been studied extensively in the context of ecological models. Moreover, this class of model has been studied in an eco-evolutionary context where mutation occurs slowly and can drive changes in both the shape of the consumption kernel and its norm [5]. The mean field equations for this model are given by:

$$\begin{aligned} \partial_t \chi &= \underbrace{\mathcal{S}_\pi \mathcal{D}_{mut}\left(\rho(c), \chi(c)\right)}_{\text{mutation}} - \underbrace{\gamma\chi(c)}_{\substack{\text{consumer}\\ \text{death}}} + \underbrace{Z_1 \int d^dy\, \psi(c-c')\chi(c)\rho(c')}_{\text{consumption+birth}} \\ \partial_t \rho &= \underbrace{g - \kappa\rho(c)}_{\substack{\text{resource supply}\\ \text{and washout}}} - \underbrace{Z_1 \int d^dc'\, \psi(c'-c)\chi(c')\rho(c)}_{\text{consumption+birth}} - \underbrace{Z_2 \int d^dc'\, \phi(c'-c)\chi(c')\rho(c)}_{\text{consumption}} \end{aligned} \tag{23}$$

Now the fixed point is given by:

$$\rho^* = \gamma/Z_1 \qquad \chi^* = \frac{gZ_1 - \kappa\gamma}{\gamma(Z_1+Z_2)}$$

with validity condition given by $\frac{gZ_1}{\kappa\gamma} > 1$ which can once again be interpreted as the condition that consumer per capita births at resource saturation ($gZ_1/\kappa$) are greater than per capita deaths ($\gamma$). We define $\nu \equiv g/\gamma$, $\alpha \equiv 1 - \frac{\kappa\gamma}{gZ_1}$, and $\theta(x) = \frac{Z_1\psi(x)+Z_2\phi(x)}{Z_1+Z_2}$ to obtain the dimensionless equations:

$$\begin{aligned} \partial_t \chi &= \mathcal{S}_\pi \mathcal{D}_{mut}(\rho(c), \chi(c)) - \chi(c)\left[1 - \int \psi(c-c')\rho(c')d^dc'\right] \\ \frac{1}{\nu}\partial_t \rho &= \left[1-\rho(c)\right] + \alpha\rho(c)\left[1 - \int \theta(c'-c)\chi(c')d^dc'\right] \end{aligned} \tag{24}$$

The Jacobian matrix for these dynamics is given by:

$$\mathbf{J} = \begin{pmatrix} \partial_\chi D_{mut,\chi}(\rho^*, \chi^*)\hat{\mathcal{S}}_\pi(q) & \tilde{\psi} + \partial_\rho D_{mut}(\rho^*, \chi^*)\hat{\mathcal{S}}_\pi(q) \\ -\nu\alpha\tilde{\theta} & \nu(\alpha - 1) \end{pmatrix} \tag{25}$$

For this model, the Jacobian has a similar structure to the model with self-renewing resources. The trace is always negative, while the determinant can become negative only for non-positive definite $\psi$ and $\phi$ or for replicative mutations in the limit $\alpha \to 1$ (as in the self-renewing resource case). It is interesting that $\alpha \to 1$ corresponds with the limit taken in [5, 9, 12], suggesting that replicative mutations might also drive interesting phenomenology in the context of discrete (but large) phenotype spaces.

##### 4.4 Non-translationally invariant kernels

For non-translationally invariant kernels, as is the case when we model disordered interactions (i.e. those drawn from a distribution and quenched), the analysis becomes more subtle because the linearized dynamics are no longer diagonalized in Fourier space. A version of this was scenario studied in [7] for comparable consumer and resource mutation rates with random asymmetric (but sign-constrained) interactions decorated on a hypercubic genotype space (as opposed to the Euclidean phenotype space studied here). In that work, the stochastic dynamics achieved the homogeneous state in the long term (via a Griffiths phase-like state), so long as the population size scale was large enough, the typical phenotypic difference between mutant and ancestor was sufficiently large and the mutation rate was sufficiently high to escape stochastic extinction. However, it remains to be explored whether or not strongly asymmetric mutation rates (i.e. $D_{mut,\rho} \approx 0$) can lead to (disordered) patterning, and whether or not this patterning is stable to extinctions induced by the chaotic fluctuations that are typical of disordered non-reciprocal interactions [1, 8]. Moreover, it remains to be studied how the effect of the underlying lattice topology, which might better reflect the underlying topological structure of genotype space, might impact patterning and extinction.
